## Supplemental Information for "Addressable Nanoantennas with Cleared Hotspots for Single-Molecule Detection on a Portable Smartphone Microscope"

**Supplementary Table 1. Temperature ramp used for folding DNA origami nanostructures**

| Temperature [°C] | Time [s] |
|---|---|
| 65 | 300 |
| 65 | 50 |
| 64 | 95 |
| 63 | 95 |
| 62 | 95 |
| 61 | 95 |
| 60 | 540 |
| 59 | 540 |
| 58 | 1140 |
| 57 | 1740 |
| 56 | 2340 |
| 55 | 2940 |
| 54 | 3540 |
| 53 | 3540 |
| 52 | 3540 |
| 51 | 3540 |
| 50 | 3540 |
| 49 | 3540 |
| 48 | 3540 |
| 47 | 3540 |
| 46 | 3540 |
| 45 | 3540 |
| 44 | 2940 |
| 43 | 2340 |
| 42 | 1740 |
| 41 | 1140 |
| 40 | 1140 |
| 39 | 1140 |
| 38 | 540 |
| 37 | 540 |
| 36 | 290 |
| 35 | 290 |
| 34 | 290 |
| 33 | 290 |
| 32 | 290 |
| 31 | 290 |
| 30 | 290 |
| 29 | 50 |
| 28 | 50 |
| 27 | 50 |
| 26 | 50 |
| 25 | 50 |

**Supplementary Table 2. Unmodified staple strands used to fold the DNA origami nanostructures**

| Name | Sequence (5'→3') |
|---|---|
| 1 | TTTAAATGTTTGCTGAGATTTAGGACCCACGCGAA |
| 2 | TTAGAACGCAATTAAGACAAATACATACATAAA |
| 3 | TTTAAGCAAATTCACAAAGTATTAAGAGGCTCGGA |
| 4 | TAAATACCCGGATATCATCAACGGTCAATCATAAGACCATCGATAC |
| 5 | GAAGGGATAGCGAGATAGTTCCGGCCAGGAAGAAGAATGAGGT |
| 6 | GCAACTGGCGAAAGGGGAGTAAAGTTGCCGGAGTGAGACCGGTCCAAAC |
| 7 | ACGAGGAGAGGCGGTTTGATGGTGGGGCCCACCCT |
| 8 | CGGTGTACAGACCAACAAAGCTAACGGAAAAAATCTACG |
| 9 | AATATCGGCACGCGCGGGCCGGAAGCATAAAAGCT |
| 10 | CAGAACAATATATCGGCCATCAAACACAGTTGAAAGGAA |
| 11 | TGAGGAAAACAGCCTGATTGCTTTGTTGC |
| 12 | GAACGCCTCCATATTATTTTA |
| 13 | AGTTCTGTCCCCCCCGAGGCGCTGGCAAGTGTTTG |
| 14 | CTTAAATCCCGGCGGTTGTG |
| 15 | AGCAATACTTCATCACGCAAATATCGCCAGTA |
| 16 | TTCATTTACCATATTGCGGAACAAAGAA |
| 17 | CTACAATTTTTTTGAAGAAAAAGCTTTAAAACAGAAATAAAGAAAAAT |
| 18 | CCTACATATCTAAAGCATCACCTCAAATTTGC |
| 19 | GGTGGCTCCAACGGCATTTCGCACTCAATCCACGCCATCCA |
| 20 | CGGAATTACCGTGTCGCAAGACAAAGAAAACAGTAAACAAAC |
| 21 | TTTCAATGATAAATTAATGC |
| 22 | GTCGAGGACCCGCCGCACCTTTTACATCCGCTGAGCAT |
| 23 | GTAATCAGAAACGAGCCTTTAGTGCCTTCTCAGAACGA |
| 24 | GCGACCCACCAAGTAGAATCATTAAAGGTGAAAATA |
| 25 | GTCTGAGCAAAAGAAGATAATGGGAAGGAG |
| 26 | TCACGCGTGGGAACAAATGTCACTGCGCGCCGCGG |
| 27 | ATTAGAGCATTTTTGCGAGCTGAAAAGGTCTA |
| 28 | TGTGATAAATTTAGCCGGAACGAGATATATTCTCA |
| 29 | TCCCGGGCGAAAGCCACCGTCTTTCCAGAGCCGAA |
| 30 | AATAAACCAGAATCTTTTCATAATCAGGA |
| 31 | CAGACCAGTTACAAAATAAAGGCTTCAGTAGGAGTATTATTAATGC |
| 32 | CGTAGGCGCATAACTGACCAACTTTGTTGCGCGATACATTGCAAAAG |
| 33 | AATAATAACCGGCGCAGAGAGTAATCTCGCCT |
| 34 | CATTATATTTTATCTTCTGACCTAAAGATGATCAATATA |
| 35 | AGGACGTTAAGAACGGTTTAATTTCAACGAGAAACCAA |
| 36 | AGGAGGCTTTAACGCCAAACGAACTGCTCAT |
| 37 | ACCACCCTTAGATGAGTGACCTGTCGTGCCAGAAT |
| 38 | GGTGATAAGAACTGGCATGATAATAACAGCCCTTTAATATC |
| 39 | CCCCTTTTCTTGTGTGAAATTGTTAAAGCACTTGT |
| 40 | CATTTAAACTCCATATAGATTCATCAGTGAACAAGAAACTCATC |
| 41 | AACAGACAATAGTTTATCCGCTGGTAAATGTGCAG |
| 42 | CGGATCGGATGTGCTGCAAGGCGATCAGTGCCAGGTGGAGCC |
| 43 | CCGAGCTCGAACTTGACGAAAGGTAAGAGGCATTTATTT |

| | |
|---|---|
| 44 | TGGGCACTAAAAAAGAGTCTGTCCTTTGATTTCAAACTTAC |
| 45 | GAGTCAACTAATTTAGGCAAGTAATCCTGAACAGA |
| 46 | AGAGTTCGTAAAGCTGATCTCATAAGGATTGACTGCCAGTTTGAGGCAG |
| 47 | TACGCGGGATACGAGGGCAACGGAATTATACCAAG |
| 48 | ATCCTTTGCAACAGGAAAAACGCT |
| 49 | GAAGGTATTATCACCCAGCAAAATCACCTTACCATTAGC |
| 50 | TTGCAAAGACAAAAGGGAATGAAATAGCAAGCAGCACC |
| 51 | GCAAGACTGGATAGCGTGAATCCCCTGTATGCGC |
| 52 | AGCACCCTCAAATCCTCCAGGAAGGGTCATTCCTTTAATTGTACAGGTG |
| 53 | TTTGCGTATTGACAATTCCACACAAAATTGGG |
| 54 | AAACGGACGACGTCGGTGACGCAACAGCGAGTATAGTTATTTTGATGGGG |
| 55 | ATATAATACACGTACTACACCAGCTAACACCATTCACCAGTCACA |
| 56 | TATTTTAACCTCAAAAGCTGCATTGCCTGGGGTGCCTAAATCCTTAGAC |
| 57 | AAAGGAAGCTTGATGTTGAAACCTG |
| 58 | GTCAGACCTCAAGAGAAGGAT |
| 59 | TTATCAGCTTGCTTACACTAT |
| 60 | AAAAATTAAAGCCTATTATTCTGAAGTTGATAGATTGCAAACCCTC |
| 61 | TTTGCGGGCCTCTGTGGTGCT |
| 62 | CACCGGAATCATTTCAAAATTATTT |
| 63 | TAAAGGAAGCTCTGGAACTGCGAACGAGTAGGCATAAACTGTAATGTCA |
| 64 | GAGCGTCCACTACCTCCGTAATTTTAGTTACAAAATCGCCGT |
| 65 | TACCAGAATCAAGTTTGCCTTATTTAAAAACTAATAAGACCGCCATGC |
| 66 | GCAGCAGAGGTCGTCGCAATTGCG |
| 67 | TGAGATCGGCTATAATATACCGACAGGGAAAGAGCGAAAGGAGCGGCAGT |
| 68 | CTTGGTAACGCCAGGGTACGACGTGGAT |
| 69 | CGCGCAGTATATTCGACAATGAATATACAGTA |
| 70 | AAGAGGTAGTACCTTGAGAAAGGCCGGACAATGCCATAGTAG |
| 71 | TGCACGACAATTGCGAATGCCCCCTCGGCTGGCCA |
| 72 | GCTTTGAGGACTAATACGAAGAAAACGAAAGAGGCCCCAGCGGATT |
| 73 | ATATAAAATTCATATGGTTTATTACCGAGGAA |
| 74 | GCAGTTGGTAAAAAGGCGGCCGCGTGGTGGGTGGTAGCAGGCTGCA |
| 75 | GTCCTTTCATGCATGTCCCAGTAAAGTGCCCGTATAAAAGGAGGTAATC |
| 76 | ACATTACAAAGGATTAAGGTGCCGTCGAGAGGACATGAAACAA |
| 77 | TAGTACTAAAGTACGGTGCCGAAAGATTTTTGATTGTAATTTTGTTGGGT |
| 78 | AGTGAATTTTCCTCAAACCCTCAGAGCCACCGAACCCACAC |
| 79 | TTATTCGGTCGGGTATTAGCCGTTTTTTCGATTTA |
| 80 | TCATCGTAACATTCCAAGAACATAGCCCCCT |
| 81 | GCCGCTACCACCACTGCCGTATCCGCTCGGCGCCAGCTGGTC |
| 82 | ACAGTGCTTTACCGAACGAACTGGTTGCTAGCGGTAAC |
| 83 | TGCCCGCTTTCCAGGTGTTGTTC |
| 84 | ATAGAGCCGCACTCCAAGTC |
| 85 | GCGGTCAGTATAGAAGATTAGCCCTTAAAGGGATTTTAG |
| 86 | GGGGTTTATATCGCATATGCATTGACCATTAGATA |
| 87 | ATTCTAGCGATGTGTAAAAATGAATCGGCCAAAAA |
| 88 | AAGTTTTGACGCTCAAATCCGGTATTCTAATAA |
| 89 | TACTGTGTCGAAATCCGCAAAGTATAGCAAC |

| | |
|---|---|
| 90 | TATTAAATCATACAAAATCATAGCGTCAAATTAT |
| 91 | CACGGGGGTAATAGTAAAACAGTTAGACGTTAGCCCTCAACAACCCAG |
| 92 | GACACGTAGATCCTTATTACG |
| 93 | ACCAACATGGCGCGTAACGATCTTACAACATTTTG |
| 94 | TTAAAGAGATCTATGACCGCTAAATCGGTTGTCCC |
| 95 | AAAAGAATTTCTTAAACATTACGAGACCAAAA |
| 96 | CCTAGTTTCCTTTCACCACTTGTAGCAGCACCGACAGTATCGGCCTACCG |
| 97 | CTGTCATACCGGCCCTGGCCCTGAGAAGA |
| 98 | AACTGTAAAACGACGGCTAAGTTGCGC |
| 99 | AAAGTCTTTCCTTATAAGAGTGTACACAGACAGTAAATGAG |
| 100 | GCAAACCACGGTTTTGTCACAATCAAAAGTAACCG |
| 101 | CATTGAAGACAGTTCATGAGGAAGTTGGGTAAATAC |
| 102 | AATTGTTTCATTCCATATTCAAAAAGCTATCAATTG |
| 103 | AGAGAGAAATAACAAGCGTTTGCCATAAGTA |
| 104 | TCAATGCTCAGTACCAGGGAGACTCGATTGGCCCA |
| 105 | ACCTTATGCGATTTTGGGAAGACAACATTAA |
| 106 | TAGTATCAAATTCTTACAGGCGTTTTAGCGAAACG |
| 107 | AGCGGGAGCTAAACAGGAGTTTTTACAATAGATTT |
| 108 | ACGGAGCCGTTAATCAGTGAGGCCTTG |
| 109 | TTTGACCGCCAGGAAAGCTAATCAGAGCAAACAAA |
| 110 | AGGAAGCGCAGCGATCCCGTGCCGCCGGAACGTAAACGATGCTGATACG |
| 111 | AGGACGTCAGACTGTAGC |
| 112 | ACTGTATCACCGTACTCCAGTTAACTGAATTCCGCCACTACGTGAAAATC |
| 113 | GAAAATTCGCAGGCGCTCAGATGCCGGGTTAATCTCCAAAGAGAACCTG |
| 114 | TCGCCGGCTGGAGGTTTCTTTGCTCACTTTTGGGTAGCTACT |
| 115 | CGACACGCCAAATTACCGCGCCCAAAATCCAAGCC |
| 116 | CAGAGCGGGGTCATTGCGTCTGGCCGGTTGAGCAGTCTTGCCCCC |
| 117 | TCCCATGCGTTCTTTGCCGATTTTCAGGTTTACGG |
| 118 | TAAAAGGAATGGCTATTAGTCGAACTGAAAAA |
| 119 | TCAGTGAGAATCAAATCAGATATAGAACAGCCCTCAGAGTACCGTTAATC |
| 120 | CTATGAGTAATGTGTAGAAAAGGGTTAA |
| 121 | AGACCGGCAAACGCGGTCCGTTTT |
| 122 | GGACAAATCACCTCAATATGAAAATTTGACGCTCA |
| 123 | TTTGACCAAAAGAAATACGTAATGCCACAGACTTTCATC |
| 124 | AAAAATAGGAGCCGGGCTCAGCAAATCGTTAAAAGGAGGCC |
| 125 | AATCAAGAATTGAGTTAAATAGCATTTTTTGTTATCCCTAGCAAGCGCC |
| 126 | GAATTGCCAGAATTCAACTATTACACCCAAATACCAGAACGAGTAG |
| 127 | GTTGCGTCGGATTCTCGTAGCATTCCTCGTAA |
| 128 | AGCCAACGTGGCACCAGAATCTTACCAACGCTACC |
| 129 | GCCACGAAACGTTCGCCACGTGCATCCGTAATGGGATAGGGCC |
| 130 | ATCCTGAAAACAAACCTTTTTTAATGGACGCGAGAGGTTTGA |
| 131 | TGCCTATAATAGGTATTATAGGATAAAAGCATAGTAAGAGCATCGA |
| 132 | ATCAAGATTGTTTGTATTCCTGATTATCATTTAATAAACTTT |
| 133 | CAAGGGGCAACTCATGGTCATAGCTAAGGGAGAGA |
| 134 | ACCGAGGCTGGCTGACCTTTCATTAGGTAGAAACCAGTC |
| 135 | GAGAACAAGCAAAACCAAATCAATATTTCGTCACTACAAGGATTTT |

| | |
|---|---|
| 136 | TTTGGACATTCTGGCCAATTGGCAGGCCTGCA |
| 137 | TGTACGGAGGGAAGTGAGCGCTTTAAGAATAGAAAAGAAACGCAAA |
| 138 | TACGTATCATGACTTGCGGGAGGTATCCTGAACCACCACTTGATATAT |
| 139 | ACGGAACGTCATTTAGTGATGAAGGCATAAAACTGGTGCCCCGGAA |
| 140 | GCAGCAACAATATCGAAGAACAGTAATAACATCACACC |
| 141 | GAGGGAATCCTGAGAAGTGGCCGATAAAACATATT |
| 142 | AAAACCGCCACCCTCAGATTTTAACGATACAGTCACCGGGATA |
| 143 | GTTTACCAGACGACTCAGAAGAGTCTGGAAAAGCCCAAA |
| 144 | AGACAATCGCCATTAAAAAAGAATCAGCAGA |
| 145 | TAGCGAGTCTTTACTCGATGATGTACCCCTTCCTGCTG |
| 146 | ATAACGGTAATTTTCACACCGATAGAAAGAG |
| 147 | TTCAAATTGAATTAATTAATT |
| 148 | GTACGAACGTTATTAATCTGTTTACTTTTTAATTAAAGCGA |
| 149 | TGTGCGGTTGCGGTATGCTCA |
| 150 | AGGCTTGCCCTGACTTTAATC |
| 151 | TGCTTCTGTAAACGAATTA |
| 152 | ATCTAGCCAGCAGCATCCCAGCGGTGCCGGTAATAATTTCGTAAA |
| 153 | AAGTTTGACCATAACAAAGTTTTGTCGAAGGAATGACAACAGGA |
| 154 | GGACGTCACCCGGTCGCAGTTTCATGTGCACGTTT |
| 155 | AATCAAATTAGTACCGCCACCGAGTAACGCGTCATCCGGAACCGCGCCTAAC |
| 156 | CGGAGAGCGGGAGAAATAAAGCCTCAGAATT |
| 157 | ACAGTGCGACTTTACAAACAAAAGCCAAGTCAATACTATCATTTCC |
| 158 | TACATCAAACTGAAAAAGAGACGCATACCAGTCGG |
| 159 | CGTGTGAATTATTAAGAGGGAGAAACAATAAACGTCAGACTCG |
| 160 | ACTAAATGGGCTTGAGATTGGCT |
| 161 | TGAGCAAAGCGTAAGTATAGCCCGGTTCGGAACCAGAATCCCTCAGAAAC |
| 162 | TCACAGAGAGTAACCCAAGCTATCCCAGCGCACGGAAATTGCAAC |
| 163 | ATACAGAACCCTTCTGACGTCTGAAAGAGCCA |
| 164 | GATAAAATCAGAGCCGGGACATCCCTTACACTAAA |
| 165 | CGCCAGCCAGAAAGCGTACTGAGTATGGTGCT |
| 166 | ATCCATGTAATAGATTAAGCACGTATAACGTGCGCTAGTTT |
| 167 | CATAACAGTTGATTACTCGGT |
| 168 | AACAAAATCGGCACGCTGCGCGTAACAGGGCGTTT |
| 169 | TGAAAGCCCAAAAGAAACCGACATTAGGGAGG |
| 170 | CCAGAGCGCCATACAGCGCCATGTTGATTCAGAAGCTAACAG |
| 171 | TTCCTCGCACGCTGATGGATTATTTACACAGAGATGTGGCAC |
| 172 | CTTAGCATCAGACGATCCACAACTATCTTTCCCAG |
| 173 | TACGCCAATTTAGAGCTTAATCTCACCCACCATAAGAAA |
| 174 | TATTTGCCGTTGCACATCTGCCCTTCACCGGTGTA |
| 175 | ACCATCGATAGGCCGGAAATTAGAGCGTCACCGACT |
| 176 | TTTAGAACCCTCATATATTTTAAATGGACAGTCGGTCAGG |
| 177 | TAGCATTTTGGGGCGCGGATGGCTTAGATCCAACA |
| 178 | AGCAAACGCTTAATAGCTATATTTTCATAACATCCAATA |
| 179 | TAATTACTAGCCTTAAATCAAGATTTTGCACAGCATTGGAGGCAG |
| 180 | TGATCGGGAAAGCTAACTCACATTTATTAATGCTTAGGTTG |
| 181 | GAAAGGAAGGGAAGAACCGGCGATCCCCGGCCGTGAGAGCCTCCGTCACGT |

| | |
|---|---|
| 182 | GAAGGTTATCTAAAAT |
| 183 | AAGGCCGCTTTTTGCG |
| 184 | CACCCTGAACAAGCCG |
| 185 | CTCGTCGCTGGCCCTCCTCCGTGCCTTAATTTAGAAACCAGTAC |
| 186 | TTTGGAACAAGACGCCGCCCCAG |

**Supplementary Table 3. Modified staple strands used for the immobilization of the DNA origami structure (biotinX), nanoparticle binding (npbindX) and fluorescence labelling.**

| Name | Sequence (5'→3') |
|---|---|
| biotin1 | biotin - AGAATATAAAGTCCCATCCGTTCTTCGGGG |
| biotin2 | biotin - AGTTACCAGAAGGAAAGCAGATAAGTCAGAGGGTAATCGCA |
| biotin3 | biotin - ACAACTTTCAACTGAGGCTATGT |
| biotin4 | biotin - AGGGCGATCGGTGCGGTGCGCAACCGGAAACAATCGGCGGG |
| biotin5 | biotin - TTCATCGGCATTGACGGGACCAATAGACCCTCAATTCATTCCAA |
| biotin6 | biotin - TAGATGGGCGCATCGTAACTTCAGGCGCCT |
| npbind1 | CATTTCGTCAACATGTTTTAAGTTTTAATTCGAGAAAAAAAAAAAA AAAAAAAA |
| npbind2 | GGTTATATAACTATATGTGAATAAAAAAAAAAAAAAAAAAAAAAA |
| npbind3 | ACCATCAACCGTTCTAGCCGCAAAAAAAAAAAAAAAAAAAAAA |
| npbind4 | ATAAAAATGCTGATGCAATGTGAAAAAAAAAAAAAAAAAAAA |
| npbind5 | AAAGAATTAGCAAAATTAAGCAGCCTTTAAAAAAAAAAAAAAAAAAAA |
| npbind6 | ACCACCAAAGGGTTAGAACCTCAATTACGAATAACCTAAAAAAAAAAAAAAAAAAA |
| npbind7 | AATCATACAGCCTGTTTTGCTGAATATAATGCGAAAAAAAAAAAAAAAAAAAAA |
| npbind8 | AATATAATCCAATGATAAATAAGGCGTTAAAAAAAAAAAAAAAAAAAAAA |
| npbind9 | AAATCACCATCAATATGATATGACCGGAAAAAAAAAAAAAAAAAAAAA |
| npbind10 | CTTCAAAGCTGTAGCCAAATGGTCAATAAGCAAGGCATAAAAATTAAAAAAAAAAAAAAAAAAAA |
| npbind11 | AAAAGTTTGAGTAACATTATCAAAAAAAAAAAAAAAAAAAAA |
| npbind12 | AATACCGATCATCAGATTATACTTCTGAATGATGACATAAATCAAAAAAAAAAAAAAAAAAAA |
| base_dye ATTO542 | TTTGTGATCTCACGTAAATTTCTGCTCA-ATTO542 |
| hotspot_dye ATTO647N | TAATCACTGTTGCCCTGATTAAATACGTTAATA-ATTO647N |
| hotspot_dye AlexaFluor 647 | TAATCACTGTTGCCCTGATTAAATACGTTAATA-AlexaFluor647 |

**Supplementary Table 4. Modified staple strands used for the sandwich detection assay: 3 capture staples (captureX), synthetic 34 nt target strand (target34) and Alexa. Fluor 647 imager strand (Alexa647 imager). Complementary regions are depicted in the same colour. If necessary, some unmodified strands from Supplementary Table 2 and modified strands from a Supplementary Table 3 should be leave out.**

| **Name** | **Strands to leave out** | **Sequence (5'→3')** |
|---|---|---|
| capture1 | hotspot_dye strand from Table3 | TAATCACTGTTGCCCTGATTAAATACGTTAATATTTTTCGG GCAATGTAGACA |
| capture2 | 186 from Table 2 | TTCGGGCAATGTAGACATTTGGAACAAGACGCCGCCCCAG |
| capture3 | 156 from Table 2 | TTCGGGCAATGTAGACACGGAGAGCGGGAGAAATAAAGCC TCAGAATT |
| target34 | | TGTCTACATTGCCCGAAATGTCCTCATTACCATA |
| Alexa647 imager | | TATGGTAATGAGGACAT-AlexaFluor647 |

**Supplementary Table 5. Modified staple strands used for solution synthesis of NACHOS. Overhang modifications (modificationX) exchange biotinX staples from the Supplemetary Table 3 of the DNA origami structure. Complementary regions are depicted in the same colour. If necessary, some unmodified strands from Supplementary Table 2 and modified strands from a Supplementary Table 3 should be leave out.**

| **Name** | **Replacing strand** | **Sequence (5'→3')** |
|---|---|---|
| modification1 | biotin1 Table 3 | GTGATGTAGGTGGTAGAGGAAAGAATATAAAGTCCCATCCGTTCTTCGGGG |
| modification2 | biotin2 Table 3 | AGTTACCAGAAGGAAAGCAGATAAGTCAGAGGGTAATCGCA |
| modification3 | biotin3 Table 3 | GTGATGTAGGTGGTAGAGGAAACAACTTTCAACTGAGGCTATGT |
| modification4 | biotin4 Table 3 | AGGGCGATCGGTGCGGTGCGCAACCGGAAACAATCGGCGGG |
| modification5 | biotin5 Table 3 | TTCATCGGCATTGACGGGACCAATAGACCCTCAATTCATTCCAA |
| modification6 | biotin6 Table 3 | GTGATGTAGGTGGTAGAGGAATAGATGGGCGCATCGTAACTTCAGGCGCCT |
| mag1 | | TCTCCATGTCACTTCTTCCTCTACCACCTACATCACCTTCTTCTTCTTCTT - biotin |
| mag2 | | GTGATGTAGGTGGTAGAGGAA |
| mag3 | | AAGAAGAAGAAGGTGATGTAGGTGGTAGAGGAAGAAGTGACATGGAGA |

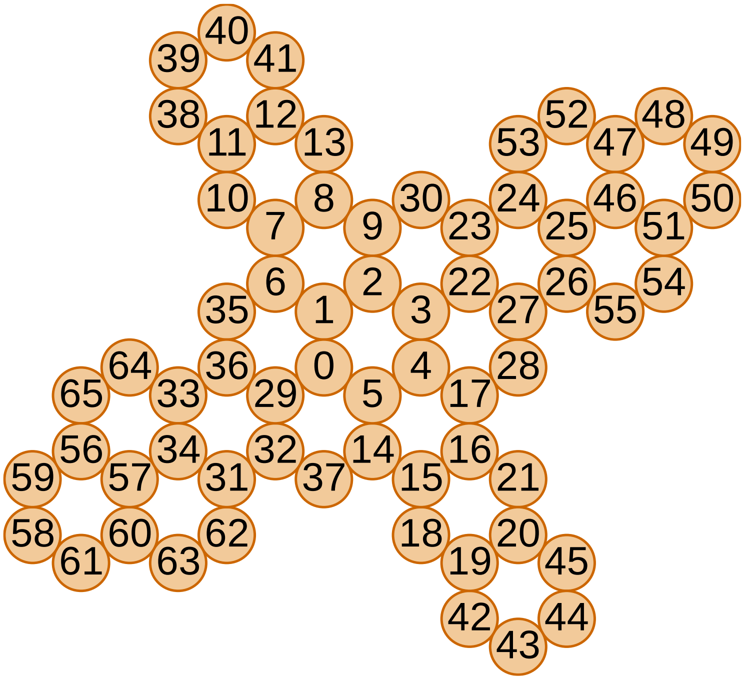

**Supplementary Figure 1. Base layout of the DNA origami nanostructure used to build NACHOS**

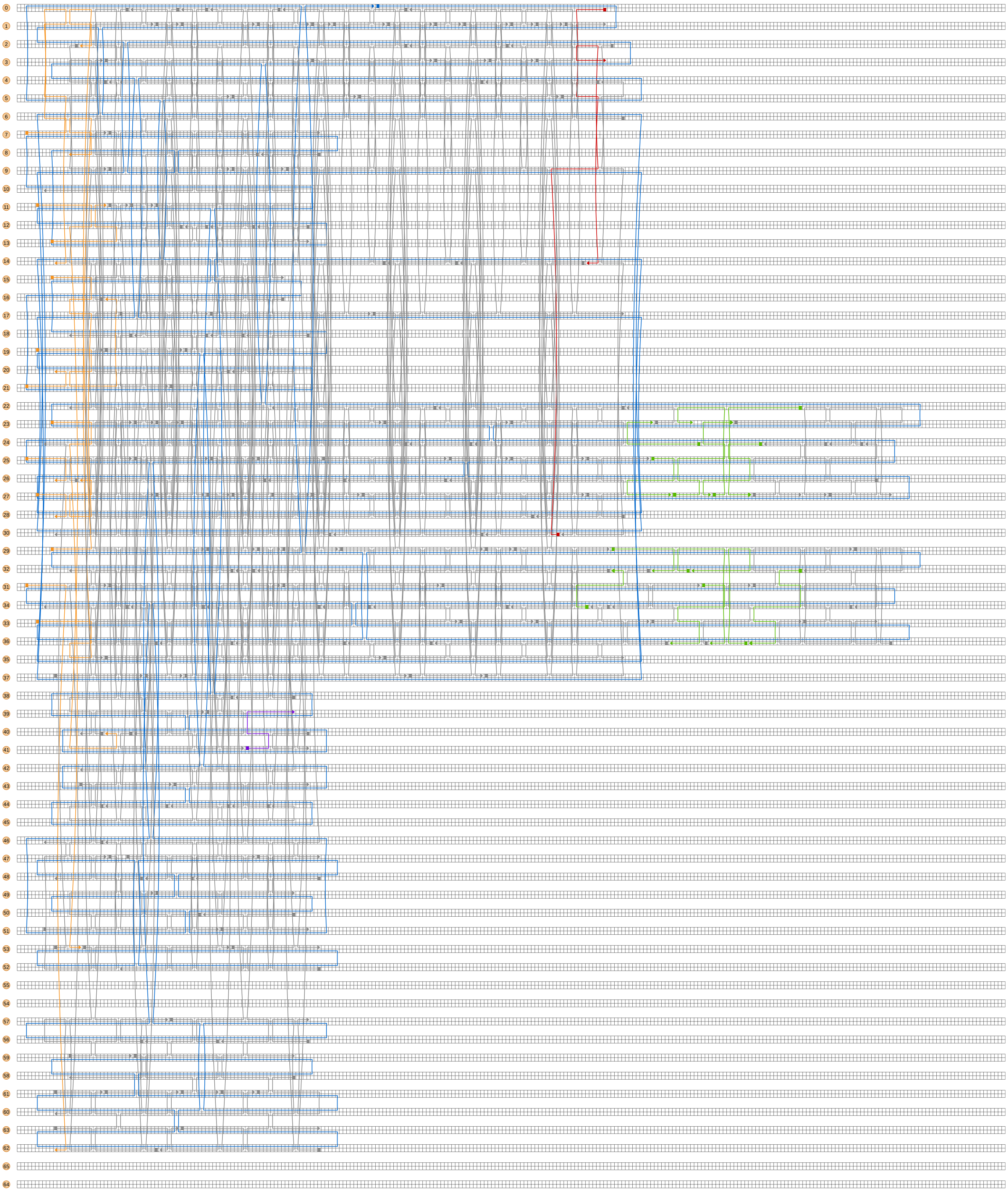

**Supplementary Figure 2. Staple layout of the DNA origami nanostructure used to build NACHOS (Yellow = biotin staples, red = hotspot staple, green= nanoparticle binding staples, purple = base dye staple)**

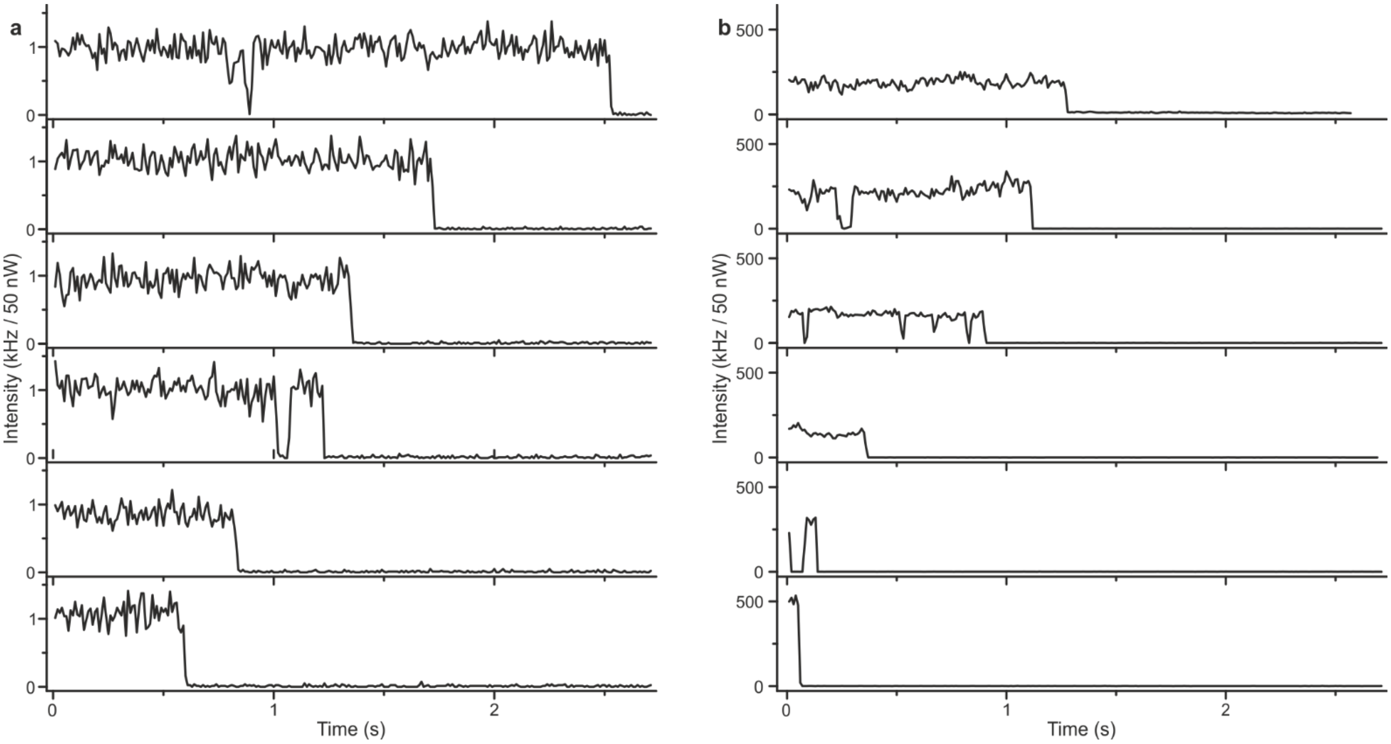

**Supplementary Figure 3. Exemplary single-molecule fluorescence transients of Alexa Fluor 647 dye in DNA origami reference structures without nanoparticles (a) and in NACHOS (b). The samples are measured at 639 nm with 200 nW and 50 nW excitation power for panel (a) and (b), respectively, and the transients are normalized to the same excitation power.**

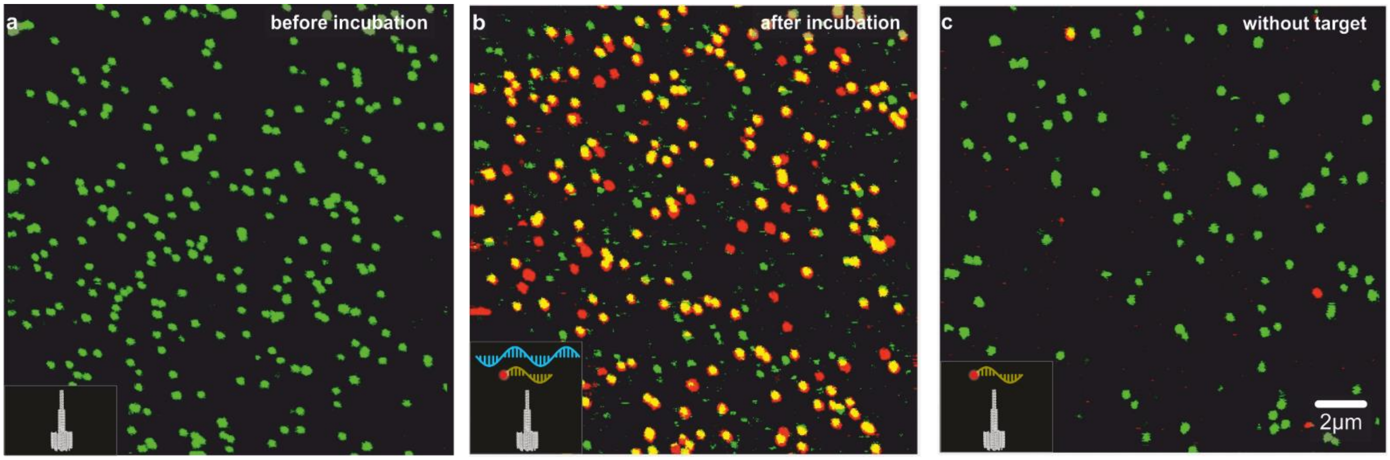

**Supplementary Figure 4. Fluorescence scans of the DNA origami reference structure (without nanoparticles) acquired before incubation (a), after incubation with the full sandwich assay (b), and after incubation with the imager strand alone. Measured at 532 nm and 639 nm with 2 μW excitation power.**

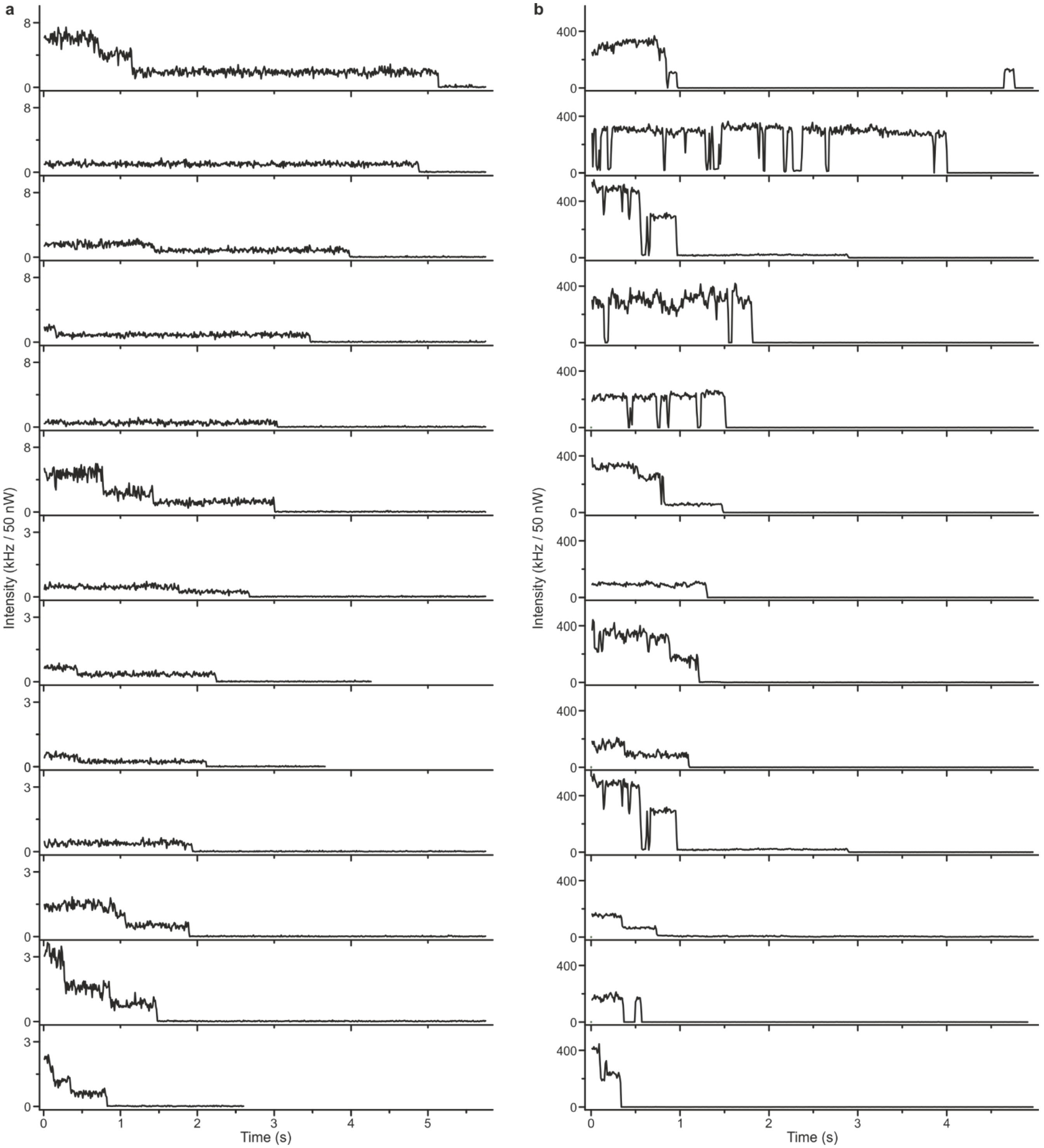

**Supplementary Figure 5. Exemplary fluorescence transients of the sandwich assay in a DNA origami reference structures without nanoparticles (a) and in NACHOS (b) The samples are measured at 639 nm with 500 nW and 50 nW excitation power for panel (a) and (b), respectively, and the transients are normalized to the same excitation power.**

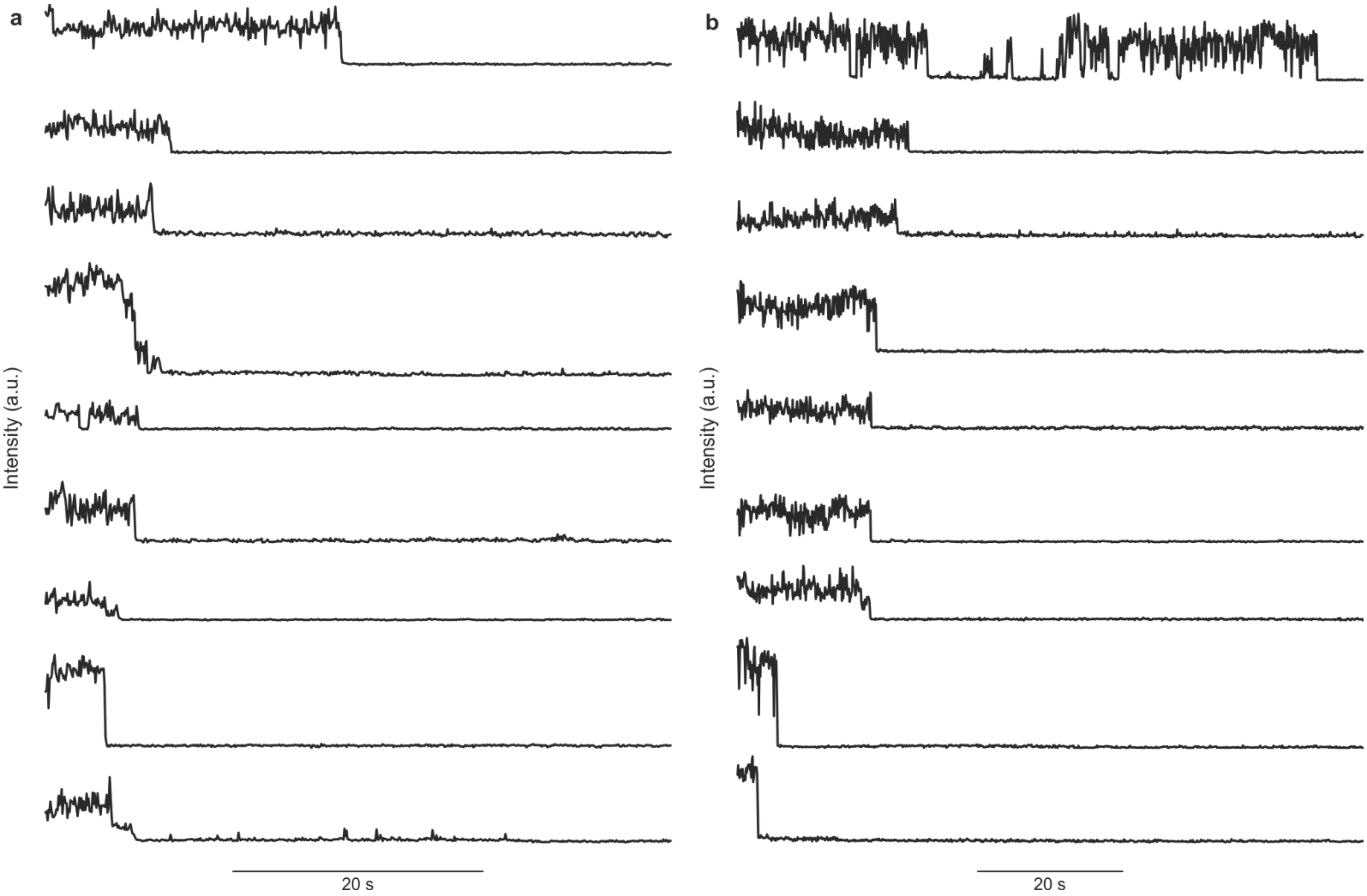

**Supplementary Figure 6. Additional fluorescence transients of single Alexa Fluor 647 dyes in NACHOS obtained from two more videos (a, b) recorded on the smartphone microscope.**

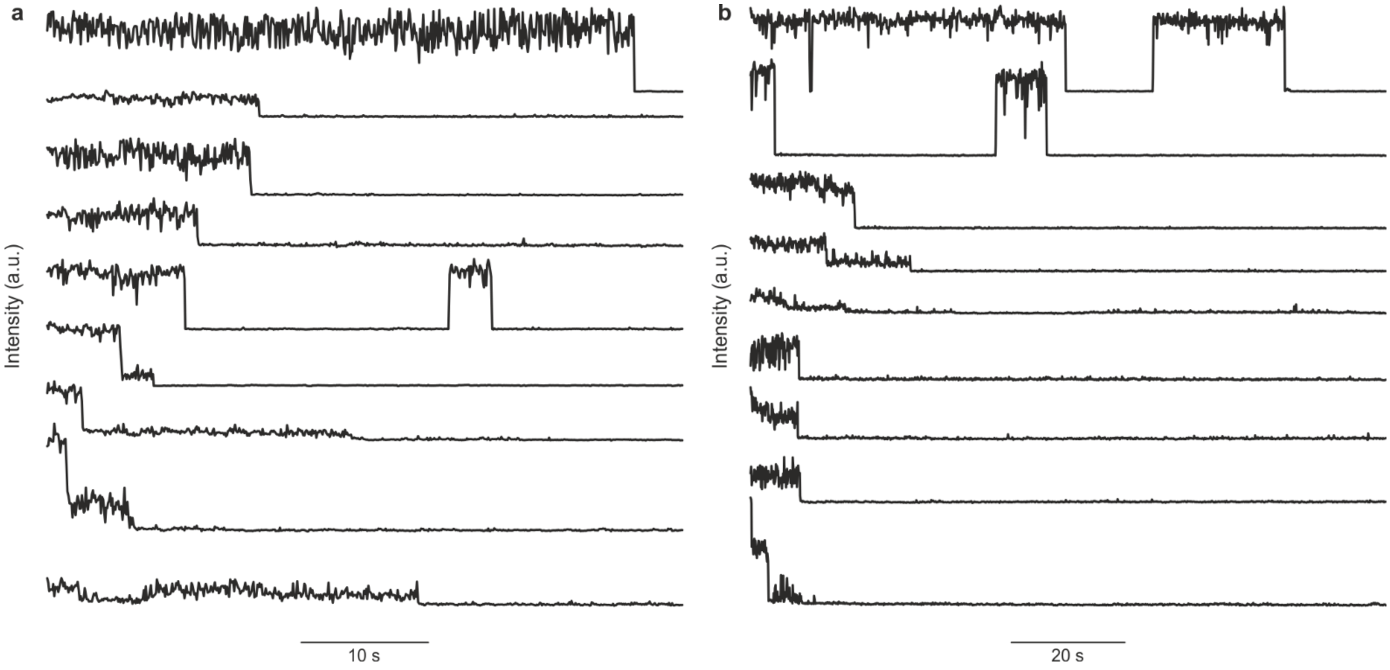

**Supplementary Figure 7. Additional fluorescence transients of the sandwich assay inside NACHOS obtained from two more videos (a, b) recorded on the smartphone microscope.**

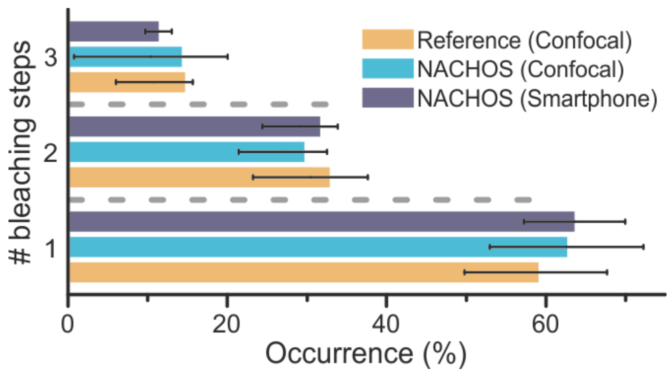

**Supplementary Figure 8. Bleaching step analysis obtained for the reference structure (orange) and for NACHOS measured on the confocal setup (blue) (same data as shown in Fig. 2g) and for 244 traces extracted from the smartphone microscope (grey). Error bars represent standard deviation from the mean.**

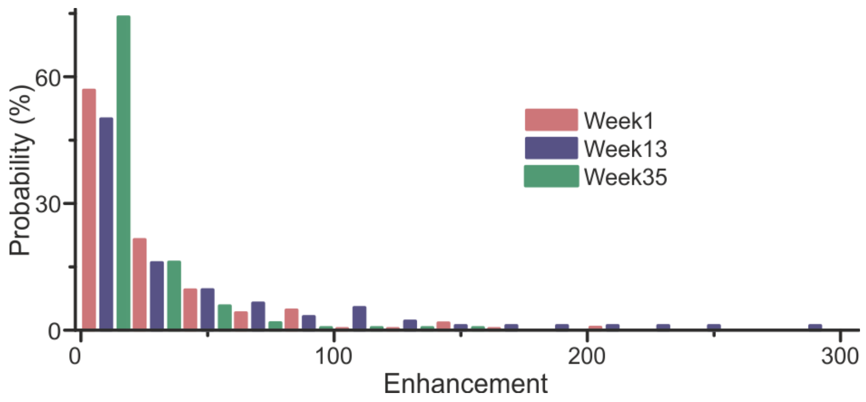

**Supplementary Figure 9. Fluorescence enhancement histograms of a single ATTO 647N dye in NACHOS of a previous design (only eight binding strands of $A_{25}$ for nanoparticles, $T_{25}$-SH used for nanoparticles functionalization). No difference between the fresh sample (red, 294 molecules measured) and the sample measured after 13 weeks (blue, 94 molecules) were observed. Slight changes are visible for the sample measured after ~35 weeks (green, 174 molecules). The sample (Lab-Tek™ II-chambers with TE buffer containing 14 mM $MgCl_2$ were stored at4 °C and care was taking to avoid drying of the sample.**

**Supplementary Note 1. Discussion pertaining the costs of the smartphone microscope**

**Price list of the current smartphone microscope**

| Name of the component | | Price |
|---|---|---|
| Excitation source: Integrated Optics 0638L-11A (Lithuania) Laser incl. power bank and cooling system | | 1892 € |
| Smartphone: Huawei P20 (China) | | 439 € |
| Objective Lens: UCTRONICS LS-40166 (USA) | | ~8 € |
| Filter: Semrock Inc. BrightLine HC 731/137 (USA) | | 472 € |
| Focussing lens: Thorlabs Inc. AC254-050-A-ML (USA) | | 114 € |
| Sample positioner: 3× Thorlabs Inc. MT1/M (USA) | | 3× 341 € = 1023 € |
| Laser positioning: Thorlabs Inc. Optomechanical Components | | 218 € |
| | Sum | ~4200 € |

**Estimated pricelist of future smartphone microscopes**

X and Y positioners can be omitted or substituted by cheaper ones since the accuracy is not needed inside the microscope.
Large scale production of the filters and adapted filter size can reduce the price by at least one order of magnitude.
Focussing lens does not have to be an achromatic one, i.e. price reduction to ~30 % of original price possible.
Smartphone can be cheaper especially if the smartphone is specialized for camera performance -> price reduction ~50 % possible. We also note that the current smartphone was purchased in early 2019 and the current value of the same smartphone is substantially lower right now. The power density in the current configuration is set to ~ 600 $\mu Wcm^{-1}$. Due to the high signal-to-background ratio we estimate that a lower power density would also be enough to make NACHOS visible on the smartphone microscope. This can be easily achieved by a high-power LED and an excitation filter to narrow down the excitation spectrum. This kind of LED in combination with a high-end excitation filter in suitable size can reduce the price to ~ 200 €.

| Name of the component | | Estimated price |
|---|---|---|
| Excitation source: e.g. Mouser, 897-LZ110R1020000 incl. power bank and bandpass filter Chroma 620/60 ET (USA) | | 200 € |
| Smartphone | | 220 € |
| Objective Lens: UCTRONICS LS-40166 (USA) | | ~8 € |
| Filter: Semrock Inc. BrightLine HC 731/137 (USA) | | 45 € |
| Focussing lens | | 37 € |
| Sample positioner (Z axis): Thorlabs Inc. MT1/M (USA) | | 341 € |
| Laser positioning: Thorlabs Inc. Optomechanical Components | | 218 € |
| | Sum | ~1000 € |

Additional discounts of at least 30 % can be expected for large scale purchase of the single components -> final price < 700 € possible.
